# Breathing strategies to influence perception: Evidence for interoceptive and exteroceptive active sensing

**DOI:** 10.1101/2025.05.15.654033

**Authors:** Francesca della Penna, Andrea Zaccaro, Başak Bayram, Francesco Bubbico, Mauro Gianni Perrucci, Marcello Costantini, Francesca Ferri

## Abstract

Recent research indicates that humans continuously and automatically modulate their breathing to temporally align exteroceptive stimuli with specific phases of the respiratory cycle. This process has been interpreted as a form of active sensing and is associated with faster responses and improved perceptual accuracy. While converging evidence suggests that respiration also shapes interoceptive processing at both neural and behavioural levels, it remains unclear whether individuals actively adjust their breathing to optimize interoceptive performance. In this study, we examined whether healthy participants modulated their respiration during an interoceptive (heartbeat discrimination) and an exteroceptive (tactile detection) task. We analysed respiration both in terms of time-locked activity and inter-trial phase coherence relative to stimulus onset and assessed their relationship with perceptual accuracy. Our results demonstrated that participants systematically adjust their breathing in both amplitude and phase, synchronizing respiration to the anticipated (i.e., cued) onset of stimuli in both tasks. Crucially, task performance was enhanced during exhalation compared to inhalation, suggesting that respiratory modulation supports the perception of both interoceptive and exteroceptive signals.

**Significance statement:** This study reveals that humans not only synchronize their breathing to anticipated external and internal stimuli, but also perform better when perceiving them during exhalation. By showing that respiration is modulated in both interoceptive and exteroceptive contexts, our findings extend the concept of active sensing to internal bodily awareness. This has important implications for understanding the dynamic interplay between physiology and perception and may guide interventions aimed at improving clinical outcomes in conditions where interoception is disrupted.

## 1. Introduction

The central nervous system continuously receives information from the body and the external environment through distinct specialized pathways. The perception of tactile, auditory, and visual stimuli is referred to as exteroception (Kassab & Alexandre, 2015), whereas interoceptive signals encompass sensations related to cardiac, respiratory, and gastric activity, which provide insight into the body’s physiological state and play a crucial role in homeostasis and adaptive behavior (Berntson & Khalsa, 2021; Engelen et al., 2023; Mehling et al., 2013). Among these systems, recent evidence highlighted respiration as a key modulator of cognitive, emotional, and sensorimotor processes (Folschweiller & Sauer, 2021; Tort et al., 2018). Notably, inhalation enhances emotional recognition (Mizuhara & Nittono, 2023; Zelano et al., 2016), episodic memory encoding and retrieval (Arshamian et al., 2018; Zelano et al., 2016), and visuospatial perceptual accuracy (Kluger et al., 2021; Perl et al., 2019). It also reduces reaction times in affective decision-making (Brændholt et al., 2024) and responses to visual (Flexman et al., 1974) and auditory (Gallego et al., 1991) stimuli. In contrast, exhalation improves conscious tactile perception (Grund et al., 2022), responses to auditory stimuli (Mizuhara et al., 2024; Münch et al., 2019), associative learning (Waselius et al., 2019, 2022), and the onset of voluntary movement (Park et al., 2020, 2022).

Furthermore, studies have reported significant task-related respiratory phase coherence across trials, indicating that participants synchronize their breathing with the expected onset of stimuli (Grund et al., 2022; Harting et al., 2024; Huijbers et al., 2014; Johannknecht & Kayser, 2022). This alignment is thought to optimize sensory processing, enhancing reaction times and accuracy (Harting et al., 2025), and aligns with the concept of active sensing, a process where organisms actively regulate sensory input to optimize perception and cognition (Allen et al., 2023; Boyadzhieva & Kayhan, 2021; Corcoran et al., 2018; Schroeder et al., 2010). According to this perspective, due to its rhythmic nature and voluntary control (Herrero et al., 2018; Maric et al., 2020; McKay et al., 2003), breathing may serve as a mechanism for actively gathering and processing sensory information.

While the role of breathing in exteroception is well-established, its influence on interoception remains underexplored and has been primarily examined in relation to cardiac signal processing. Recent studies from our laboratory (Zaccaro et al., 2022, 2024) showed that the heartbeat-evoked potential (HEP), a neural correlate of cardiac interoception (Coll et al., 2021; Mai et al., 2018; Schandry et al., 1986), increases in amplitude during exhalation compared to inhalation while performing a cardiac interoceptive task, with higher HEP amplitude associated with improved cardiac interoceptive accuracy. These findings suggest that respiration modulates cardiac interoception, yet it remains unclear whether individuals actively adjust their breathing to enhance interoceptive processing in the same way they do for exteroception.

In this study, we investigated the relationship between respiration and the perception of anticipated (i.e., cued) interoceptive and exteroceptive stimuli. We first adapted a modified version of the heartbeat discrimination task (HDT) to allow for trial-by-trial analysis of task-related breathing modulations and their relationships with perceptual accuracy. To directly compare respiratory modulations across interoceptive and exteroceptive domains, we also included a modified tactile detection task (TDT, Grund et al., 2022). Based on previous findings (Grund et al., 2022; Zaccaro et al., 2022, 2024), we hypothesized that participants would actively modulate their breathing during both tasks, temporally aligning respiration with trial onset to optimize perceptual accuracy. Furthermore, we expected that greater respiratory phase coherence across trials would be related to higher perceptual accuracy, especially during exhalation.

## 2. Materials and methods

### 2.1. Ethics statement

The study was approved by the Ethics Committee of the Department of Psychology, “G. d’Annunzio” University of Chieti-Pescara (Protocol Number 23014). The study was conducted in accordance with the guidelines of the Italian Association of Psychology and the Declaration of Helsinki and its later amendments. Participants were unaware of the specific aims of the experiment. After reading the consent form, all participants provided written informed consent.

### 2.2. Participants

Forty-one healthy volunteers (25 females; 3 left-handed; mean age: 24 ± 4.74 years [mean ± SD]) participated in the study. Eligibility was determined based on self-reported criteria, including: (1) no personal or family history of psychiatric, neurological, or somatic disorders; (2) no chronic or acute respiratory conditions; and (3) no use of medications affecting pulmonary or cardiovascular function. All participants had a body mass index (BMI) within the normal range (24.07 ± 5.36 [mean ± SD]) and normal or corrected-to-normal vision. For the exteroceptive task (TDT), inclusion required an accuracy range of 25% to 75% and a d’ index greater than 0 (Grund et al., 2022). For the interoceptive task (HDT), participants were included if their accuracy exceeded 55% (Kleckner et al., 2015) and their d’ index was greater than 0 (Sokol-Hessner et al., 2015; Whitehead et al., 1977).

### 2.3. Apparatus and stimuli

The experiment was conducted at the Institute of Advanced Biomedical Technologies (ITAB) of the “G. d’Annunzio” University of Chieti-Pescara. Participants were recruited via flyers and social media advertisements. Electrophysiological data were recorded using a BIOPAC system (MP160, BIOPAC Systems Inc., Goleta, CA, USA) and AcqKnowledge software. Electrocardiogram (ECG) data were acquired using three Ag/AgCl pre-gelled electrodes positioned over the left and right clavicles and the right costal margin (ground electrode). Respiration was recorded via a chest-placed respiratory transducer (TSD201, BIOPAC Systems Inc., Goleta, CA, USA). Near-threshold tactile stimuli consisted of square-wave electrical pulses (0.2 ms duration) delivered through a constant current stimulator (Digitimer DS7A, Hertfordshire, UK) via electrodes attached to the left index finger (anode: medial phalanx; cathode: proximal phalanx). Stimulation intensity remained constant throughout the experiment, as determined during the initial threshold assessment (mean: 1.38 mA; range: 0.8–2.4 mA). Auditory stimuli consisted of three consecutive pure tones (1000 Hz, 30 ms duration, 60 dB) presented via a buzzer, synchronized in real-time with participants’ heartbeats as detected by the ECG. All tasks were administered using E-Prime 3.0 (Psychology Software Tools, Pittsburgh, PA, USA).

### 2.4. Experimental design

The experimental session included a cardiac interoceptive task (HDT) and a somatosensory exteroceptive task (TDT), counterbalanced across participants. The HDT was preceded by an audiovisual (AV) familiarization task (Betka et al., 2021), and the TDT was preceded by an individual somatosensory threshold assessment. The entire session lasted approximately 1.5 hours. Experimental procedures are depicted in Figure 1.

**Figure 1.**
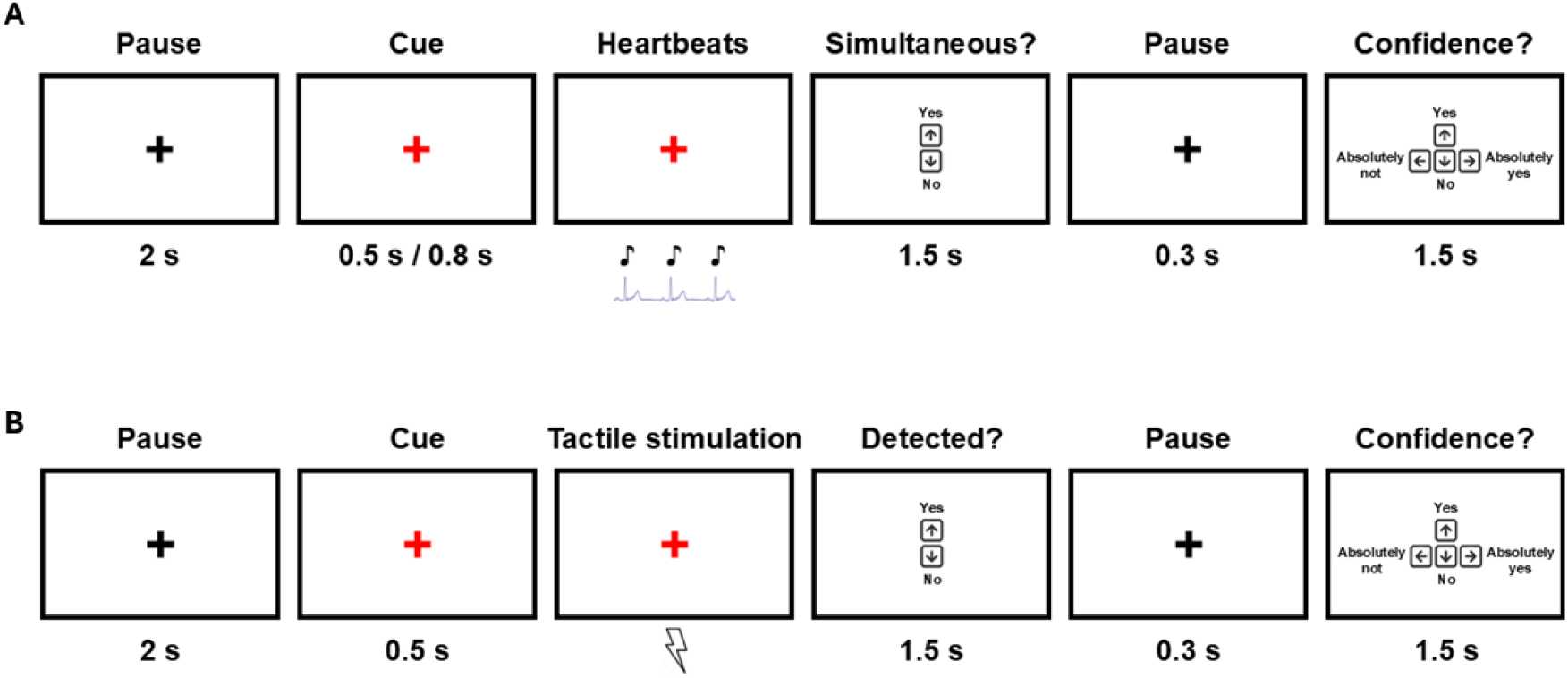
Experimental procedures. **A)** HDT: each trial began with a neutral fixation cross (2 s), followed by a cue (0.5 or 0.8 s) during which the cross turned red and the three tones were presented. After the stimulation, a simultaneity question appeared (1.5 s), followed by a fixation cross (0.3 s) and a confidence rating (1.5 s). **B)** TDT: each trial began with a neutral fixation cross (2 s), followed by a cue (0.5) after which the electrical pulse was delivered. After the stimulation, a simultaneity question appeared (1.5 s), followed by a fixation cross (0.3 s) and a confidence rating (1.5 s).

#### 2.4.1 AV familiarization task

In the AV familiarization task (Betka et al., 2021), participants judged the simultaneity of a flashing dot and a sequence of three auditory tones. The task comprised two conditions: simultaneous (S+) and non-simultaneous (S-). In the S+ condition, the dot and tones were presented without delay; in the S-condition, the tones were delayed by 550 ms. The task included 30 trials (15 S+, 15 S-) in random order. After each trial, participants indicated whether the stimuli were simultaneous and rated their confidence on a 4-point Likert scale (1 = very unconfident; 4 = very confident).

#### 2.4.2 Heartbeat discrimination task

In the HDT, participants focused on their heartbeat and judged whether it was simultaneous with three auditory tones (Betka et al., 2021; Garfinkel et al., 2015; Katkin et al., 1982; Schulz et al., 2013). Two conditions were included: S+ (tone onset delayed by 250 ms after the R-peak) and S- (tone onset delayed by 550 ms after the R-peak) (Betka et al., 2021; Garfinkel et al., 2015, 2016; Koreki et al., 2022). The task comprised 200 trials (100 S+, 100 S-), divided into four blocks of 50 trials (25 S+, 25 S- in random order). After each trial, participants provided a simultaneity judgment and rated confidence from 1 (very unconfident) to 4 (very confident). Each trial began with a 2-second white fixation cross, followed by a visual cue (fixation cross turning red). Cue duration varied by condition (S+: 800 ms; S-: 500 ms) to prevent participants from inferring the condition based on the shorter trial duration associated with the S+ condition. Following tone presentation, simultaneity and confidence ratings were collected (1.5 seconds per response; Fig. 1A). Participants were instructed not to manually check their heartbeat. Before beginning, they confirmed that the respiratory belt did not interfere with heartbeat perception. Breaks were allowed between blocks to prevent fatigue.

The HDT was adapted from Betka et al. (2021), which used five consecutive auditory stimuli. However, five consecutive heartbeats are likely distributed across more than one respiratory phase, making it difficult to assign the trial exclusively to either inhalation or exhalation. To address this issue, we first conducted a Pilot study in which HDT was presented in three different variations: 3 consecutive tones, 2 consecutive tones, and 1 single tone (see Supplemental material, Pilot study). The goal of the Pilot study was to identify the shortest possible HDT variant that would maintain the desired accuracy above 50%. The implementation of these shorter trials would allow for a precise trial-by-trial determination of respiratory phases.

#### 2.4.3 Tactile detection task

In the TDT (Grund et al., 2022), participants reported whether they perceived a near-threshold tactile stimulus on the left index finger. Two conditions were included: S+ (electrical pulse delivered) and S- (catch trials, no stimulation). The task consisted of 300 trials (200 S+, 100 S-), divided into four blocks of 75 trials (50 S+, 25 S- in random order). After each trial, participants reported whether they perceived the stimulus and rated confidence from 1 (very unconfident) to 4 (very confident). Each trial began with a 2-second white fixation cross, followed by a visual cue (fixation cross turning red) for 500 ms before the tactile stimulus. Response screens for detection and confidence lasted 1.5 seconds each (Fig. 1B). Prior to the TDT, participants underwent a tactile threshold assessment using an up-and-down method (Mylius et al., 2023): starting with a suprathreshold intensity, an initial rough estimation of the threshold was obtained by 1) decreasing the electrical intensity by 50% when the participant detected the stimulus, and 2) increasing the intensity by 50% when the participant did not detect the stimulus. A test block of 15 trials (10 stimulated, 5 catch) confirmed the threshold, defined as detection in 4–6 of 10 stimulated trials (∼50% detection rate). This procedure was repeated twice before the experiment and between each block to maintain a detection rate around 50% (never below 25% and never above 75%) throughout the task (Grund et al., 2022).

### 2.5. Cardio-respiratory data acquisition and pre-processing

Cardio-respiratory signals were recorded at a sampling rate of 2 kHz. A band-pass filter (0.1–50 Hz) and a 50-Hz notch filter were applied. The signals were then downsampled to 256 Hz. Data analysis was conducted using custom MATLAB scripts (R2023b, MathWorks Inc.). R-peaks were detected using the Pan-Tompkins algorithm (Pan & Tompkins, 1985). Respiratory phases (inhalation and exhalation) were identified using the *findpeaks* function. Inhalation onsets were marked at local minima (troughs), whereas exhalation onsets were marked at local maxima (peaks). A complete respiratory cycle was defined as the interval between two consecutive troughs. Outliers, defined as values exceeding three scaled median absolute deviations from the local median within a moving 1-second window, were linearly interpolated using neighboring values. The data were subsequently smoothed with a 1-second window filter (Savitzky & Golay, 1964) and z-scored.

### 2.6. Task accuracy assessment

Task performance in HDT and TDT was evaluated using signal detection theory (Green & Swets, 1966), which describes the ability to distinguish weak stimuli from background noise (Wickens, 2001). Within this framework, response outcomes included hits (correctly detecting the stimulus), misses (failing to detect the stimulus), correct rejections (correctly identifying the absence of a stimulus), and false alarms (incorrectly identifying the presence of a stimulus). In this study, a hit corresponded to an S+ trial accurately identified as S+, a miss was an S+ trial incorrectly marked as S−, a correct rejection was an S− trial accurately identified as S−, and a false alarm was an S− trial incorrectly marked as S+ (Forkmann et al., 2016; Katkin et al., 1982; Rominger et al., 2021; Schulz et al., 2013, 2021; Sokol-Hessner et al., 2015; Whitehead et al., 1977). This method enabled the assessment of task accuracy, sensitivity (d’), and response criterion (c).

Task accuracy was defined as the proportion of correct responses relative to the total number of trials (Garfinkel et al., 2016; Kleckner et al., 2015):

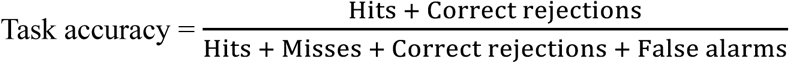

Sensitivity (d’) reflects a participant’s ability to distinguish between signal and noise (i.e., the absence of a signal). Higher d’ values indicate greater sensitivity. It is calculated using the following formula (Michal et al., 2014; Rominger et al., 2021; Schulz et al., 2021; Sokol-Hessner et al., 2015):

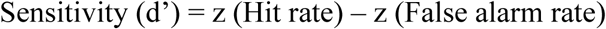

where the hit rate is the proportion of hits relative to the total number of S+ trials, the false alarm rate is the proportion of false alarms relative to the total number of S− trials, and z denotes the normal inverse cumulative distribution function.

The response criterion (c) represents a participant’s decision threshold for determining whether a signal is present. A lower criterion indicates a tendency to report the presence of a signal, leading to more hits but also more false alarms. Conversely, a higher criterion indicates a tendency to report the absence of a signal, resulting in more misses but also more correct rejections. If c equals zero, the participant is unbiased. The criterion is computed as follows (Mylius et al., 2023):

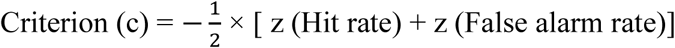

where the hit rate and false alarm rate are defined as above, and z is the normal inverse cumulative distribution function. Accuracy, d’, and c were computed separately for inhalation and exhalation. Trial classification (inhalation or exhalation) was based on the timing of stimulus presentation, specifically the first heartbeat of the trial in the HDT and the tactile stimulation in the TDT.

### 2.7. Statistical analyses

#### 2.7.1 Overview

To investigate breathing behaviour and its relationship with perceptual accuracy, we performed a series of analyses across interoceptive (HDT) and exteroceptive (TDT) tasks. First, we examined key respiratory parameters (i.e., breathing frequency, average inhalation duration, average exhalation duration) to assess differences in breathing patterns between tasks. To determine whether participants actively modulated their respiration around stimulus presentation, we computed the event-related respiratory signal (ERrS). We then assessed inter-trial phase coherence (ITC) to evaluate the consistency of respiratory phase alignment relative to stimulus onset.

To determine whether participants’ perceptual performance varied as a function of the respiratory phase at the time of stimulus presentation, we conducted paired-samples t-tests between inhalation and exhalation for accuracy, sensitivity (d’), and response criterion (c), separately for the HDT and the TDT. To examine whether respiration predicted task performance, we used generalized linear mixed-effects models (GLME). Additionally, to explore the directionality of the effect observed in the GLME, we tested for uniformity the distribution of respiratory phase angles for both correct and incorrect trials across the respiratory cycle. Finally, we explored whether respiration-related modulation of performance was driven by phase alignment (ITC), amplitude modulation (ERrS), or both, by computing ERrS and ITC for correct and incorrect trials.

#### 2.7.2 Respiratory parameters analyses

To investigate differences in breathing patterns across tasks, we calculated key respiratory parameters, including average inhalation duration, average exhalation duration, and breathing frequency, for both the HDT and TDT, using custom MATLAB scripts (R2023b, MathWorks Inc.). We assessed differences in these parameters between the interoceptive task (HDT) and the exteroceptive task (TDT) using paired t-tests.

#### 2.7.3 Event-related respiratory signal analyses

To examine whether participants actively modulated their breathing cycle and whether this modulation differed between tasks, we computed the event-related respiratory signal (ERrS) using FieldTrip (Oostenveld et al., 2011). Respiratory data were time-locked to cue onset (the white cross turning red) and segmented from -500 ms to 3000 ms. The ERrS was then calculated by averaging trials separately for the HDT and TDT. After baseline correction using a 500 ms pre-cue period (from -500 ms to 0), we analysed differences in ERrS between the two tasks. For finer-grained analysis, the 3-second active phase was divided into six 500 ms bins, and the ERrS from these bins was compared across tasks using a permutation-based t-test, with Benjamini and Yekutieli correction procedure (Benjamini & Yekutieli, 2001) applied for multiple comparisons.

#### 2.7.4 Inter-trial coherence analyses

To assess the consistency of respiratory phase alignment across trials, reflecting participants’ tendency to synchronize their breathing with trial onset, we conducted inter-trial coherence (ITC) analyses using FieldTrip (Oostenveld et al., 2011). In both tasks, the respiratory signal was time-locked to cue onset (the white cross turning red) and segmented from -500 ms to 3000 ms. The phase angle of the respiratory signal was computed using the Hilbert transform, and ITC was calculated by averaging phase angles across trials for each participant. ITC values were analysed across two distinct phases: baseline (500 ms before cue onset) and active (3 seconds following cue onset, corresponding to trial duration). For finer-grained analysis, the active phase was further divided into six consecutive 500 ms bins. To assess statistical significance, each of the six active-phase bins was compared to the baseline phase using a permutation-based t-test, with Benjamini and Yekutieli correction procedure (Benjamini & Yekutieli, 2001) applied for multiple comparisons. Additionally, ITC values were normalized for the 500 ms baseline to obtain the relative ITC (rITC), and the six 500 ms bins of the active phase were analysed across tasks (HDT vs. TDT) using a permutation-based t-test, with Benjamini and Yekutieli correction procedure (Benjamini & Yekutieli, 2001) applied for multiple comparisons.

#### 2.7.5 Behavioural analyses

Based on the computed values of task accuracy, sensitivity (d’), and response criterion (c), we performed paired-sample t-tests to assess whether participants’ performance differed between inhalation and exhalation. These analyses were conducted separately for each task (HDT and TDT).

#### 2.7.6 Generalized linear mixed-effects models

We investigated whether respiration could predict task accuracy. Behavioral data were segmented into 60 equally spaced bins based on the respiratory phase (Kluger et al., 2021, 2023; Saltafossi et al., 2025). Within each bin, we computed mean accuracy, mean confidence ratings (low confidence: responses 1 and 2 on the 4-point Likert scale; high confidence: responses 3 and 4), and the relative frequency of trials for each condition (simultaneous /non-simultaneous for HDT and stimulated/catch for TDT). This binning approach was implemented to reduce trial-by-trial variability while preserving within-subject dynamics. The aggregated values for each bin were used as predictors in a generalized linear mixed-effects model (GLME) implemented in MATLAB with the *fitglme* function. For both HDT and TDT, the first model (equations 1 and 2, respectively) employed a logistic regression framework, with task accuracy (correct vs. incorrect) as the binary dependent variable. Fixed effects included task conditions and confidence levels, while a random intercept was included for participants. A second model (equations 3 and 4) assessed whether respiration improved model fit by incorporating sine and cosine transformations of the respiratory phase angle. Phase angles were extracted via the Hilbert transform at the time points corresponding to maximum ITC values for each task. The two models were then compared using a likelihood ratio test (LRT) implemented in MATLAB.

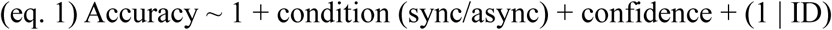

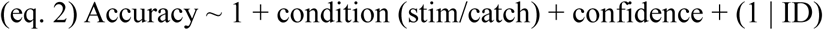

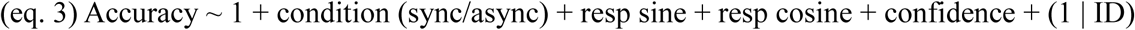

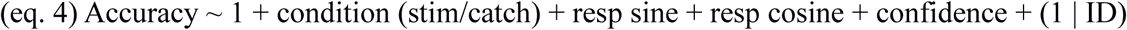

#### 2.7.7 Respiratory phase clustering analyses

To assess whether changes in task accuracy were related to the respiratory phase, we tested for uniformity the distribution of respiratory phase angles for both correct and incorrect trials across the respiratory cycle using the Rayleigh test (Berens, 2009) in MATLAB, separately for the HDT and TDT. Additionally, the Watson-Williams test (Berens, 2009) was applied to compare the mean phase directions of correct and incorrect trials within each task.

#### 2.7.8 Event-related respiratory signal analyses for correct and incorrect trials

To assess whether changes in task accuracy were related to respiratory amplitude modulation, we analysed the ERrS of correct and incorrect trials in both tasks. As described previously, the active-phase bins were compared between correct and incorrect trials within each task (HDT and TDT) using a cluster-based permutation test, with Benjamini and Yekutieli correction procedure (Benjamini & Yekutieli, 2001) applied for multiple comparisons.

#### 2.7.9 Inter-trial coherence analyses for correct and incorrect trials

We further investigated whether task accuracy was related to respiratory phase consistency across trials by analysing ITC separately for correct and incorrect trials in both tasks, segmented into baseline and active phases, as previously described. Correct trials included hits and correct rejections, while incorrect trials included misses and false alarms. To assess statistical significance, each of the six active-phase bins was compared with the baseline phase using a permutation-based t-test, conducted separately for correct and incorrect trials. To compare rITC values between correct and incorrect trials within each task, a permutation-based t-test was performed across the six active-phase bins. This methodology was applied within HDT and TDT. The Benjamini and Yekutieli correction procedure (Benjamini & Yekutieli, 2001) for multiple comparisons was applied.

### 2.8. Code accessibility

The custom codes used for data preprocessing and statistical analysis will be made publicly available upon acceptance and publication of the manuscript via a dedicated repository on GitHub. Access details, including the repository link, will be provided in the final version. The code can be made available to reviewers upon request.

## 3. Results

### 3.1. Participants selection

We evaluated 41 healthy volunteers who completed both an interoceptive and an exteroceptive task. The interoceptive task was the HDT (adapted from Betka et al., 2021), in which participants judged the simultaneity of their own heartbeat with a sequence of three acoustic signals. The choice of the three-sound condition was based on the results from the Pilot study (see Supplemental material, Pilot study). The exteroceptive task was the TDT (adapted from Grund et al., 2022), where participants detected near-threshold tactile stimuli. Thirteen participants were excluded from the HDT: six due to task accuracy below .55 (.51 ± .01 [mean ± SD]) and seven because of a d’ score below 0 (-0.06 ± .03 [mean ± SD]). This resulted in an HDT sample size of N = 28 (16 females, 3 left-handed; age: 26 ± 5.1 years [mean ± SD]). Mean accuracy was .61 ± .08 (mean ± SD), mean d’ was .68 ± .46 (mean ± SD), and mean criterion was -0.29 ± .42 (mean ± SD). Nine participants were excluded from the TDT: six due to task accuracy above .75 (.8 ± .06 [mean ± SD]) and three because of a d’ score below 0 (- 0.24 ± .03 [mean ± SD]) (Grund, et al., 2022). This resulted in a TDT sample size of N = 32 (19 females, 3 left-handed; age: 25 ± 4.7 years [mean ± SD]). Mean accuracy was .56 ± .1 (mean ± SD), mean d’ was .89 ± .63 (mean ± SD), and mean criterion was .66 ± .49 (mean ± SD).

### 3.2. Respiratory parameters analyses

To investigate breathing patterns across the two tasks, we computed the breathing frequency, average inhalation duration and average exhalation duration, for both HDT and TDT. We then conducted paired t-tests to assess task-related differences. A total of 22 participants met the inclusion criteria for both tasks and were included in this analysis. Breathing frequency was significantly lower during HDT compared to TDT (t_21_ = 3.81, p = .001, d = .81, Fig. 2A). The average inhalation duration was significantly longer during HDT than TDT, (t_21_ = -4.24, p = .0004, d = -0.9, Fig. 2B) and the average exhalation duration showed a trend for an increase during HDT compared to TDT (t_21_ = -1.87, p = .076, d = -0.4, Fig. 2C).

**Figure 2.**
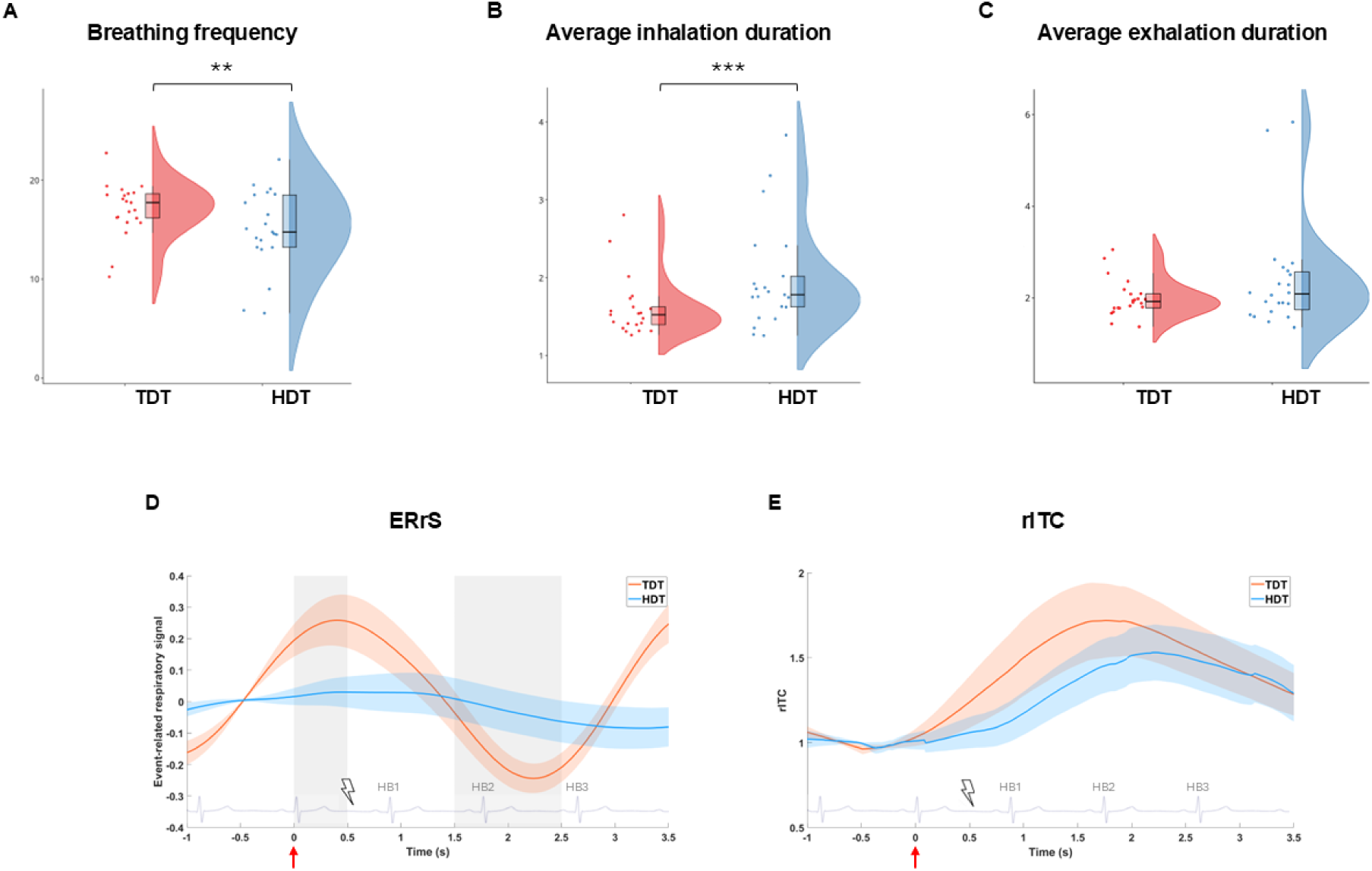
Respiratory modulations across tasks. **A)** Breathing frequency; **B)** Average inhalation duration; **C)** Average exhalation duration. **D)** ERrS. **E)** rITC. Gray areas represent the windows of statistical significance. Red arrows represent the cue onset in both conditions; lightning symbol represents the onset of the electrical stimulation in the TDT; the overlaid ECG signal represents an example of an S-trial, included to illustrate the temporal dynamics of the trial with the three target heartbeats highlighted. ** p < .01, *** p < .001.

### 3.3. Event-related respiratory signal analyses

We computed the ERrS for both the HDT and TDT to investigate if participants synchronized their breathing differently depending on task timing. A significant difference in breathing patterns between HDT and TDT was observed in two distinct time windows: 0–0.5 seconds (t = 2.56, p = .004, p_FDR_ = .022, d = .66) and 2–2.5 seconds after cue onset (t = -2.38, p = .007, p_FDR_ = .022, d = -0.61, Fig. 2D). To ensure that the observed effects were not simply due to the presentation of three consecutive stimuli during HDT, as opposed to a single stimulus in the TDT, we examined ERrS in the 1-tone condition of the Pilot study and compared it to ERrS measured during the TDT. This analysis revealed no relationship between ERrS and the number of HDT tones presented (see Supplemental material, Pilot study, Fig. S1).

### 3.4. Inter-trial coherence analyses

To assess the consistency of respiratory phase alignment across trials, we conducted ITC analyses, which revealed significant modulation in both the HDT and TDT, indicating that participants synchronized their breathing with trial onset. In the HDT, ITC values significantly increased relative to baseline within a 1.5–3 second window after cue onset (t = 2.4, p = .009, p_FDR_ = .02, d = .45). In the TDT, ITC values significantly increased within a 1–3 second window after cue onset (t = 4.39, p = .0002, p_FDR_ = .0003, d = .78). Peak ITC values were observed 2.25 seconds after cue onset during the HDT and 1.95 seconds after cue onset during the TDT. The comparison between relative ITC values in the HDT and TDT was not significant (t = .16, p = .44, d = .04, Fig. 2E), indicating that participants synchronized their breathing with trial onset similarly in both tasks (Fig. 2E).

### 3.5. Heartbeat discrimination task and respiration: accuracy, sensitivity and criterion

HDT accuracy was significantly higher for trials initiated during the exhalation phase compared to the inhalation phase (t_27_ = -2.09, p = .046, d = -0.4, Fig. 3A). While sensitivity (d’) and criterion (c) did not show statistically significant differences between inhalation and exhalation, both measures exhibited a trend towards significance. Specifically, sensitivity tended to be higher during exhalation than inhalation (t_27_ = -1.78, p = .086, d = -0.34), whereas criterion tended to be lower during exhalation (t_27_ = 1.86, p = .073, d = .35). These trends appear to be driven by differences in the hit rate, which was significantly higher during exhalation compared to inhalation (t_27_ = -2.54, p = .017, d = -0.48, Fig. 3A). In contrast, no significant difference was found in the false alarm rate (t_27_ = -.263, p = .8, d = -0.05).

**Figure 3.**
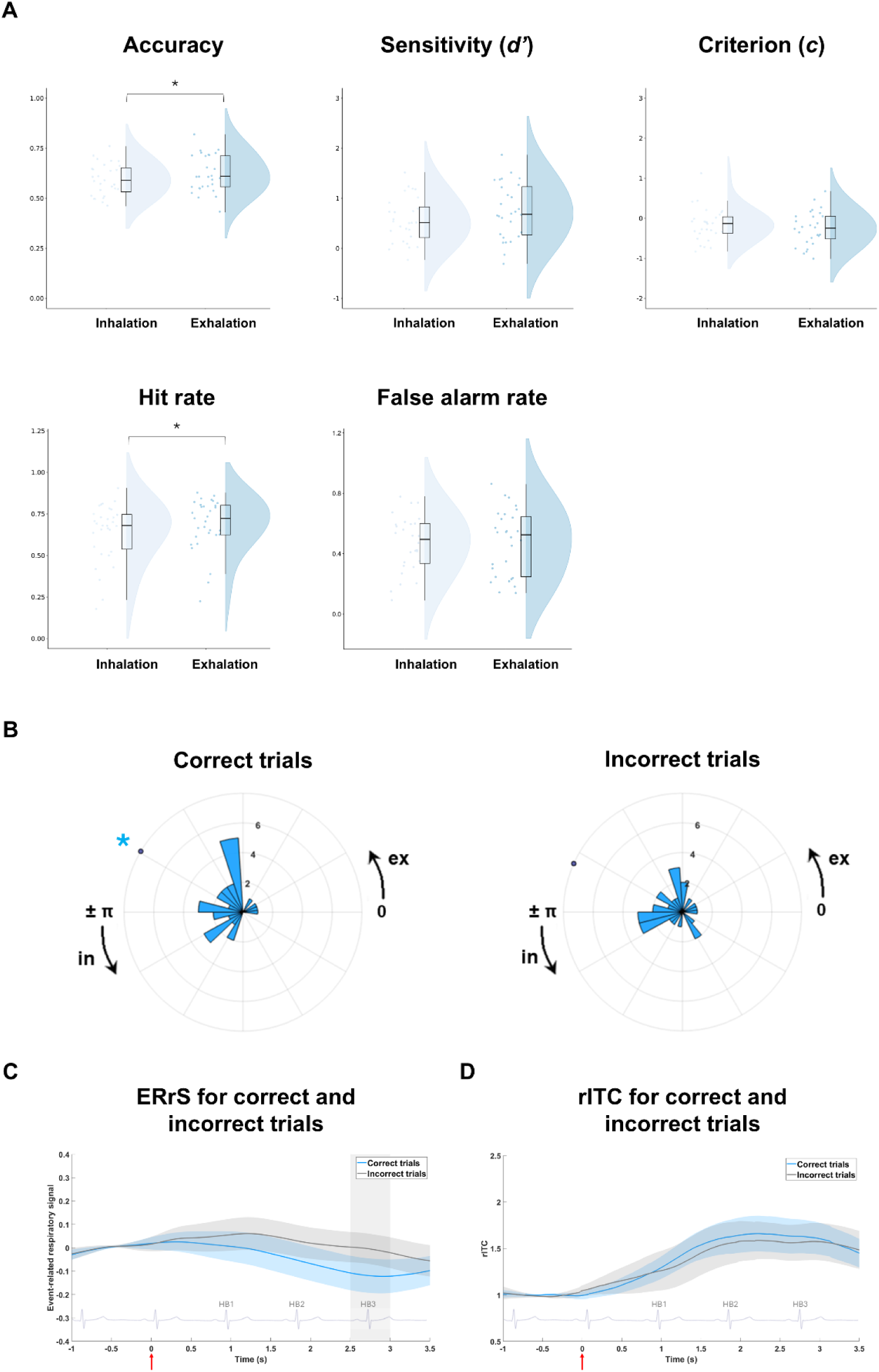
Relationship between respiration and accuracy in the HDT. **A)** Accuracy, sensitivity (d’), criterion (c), hit rate and false alarm rate obtained by dividing the respiratory signal into inhalation and exhalation phases. * p < .05. **B)** Distribution of correct (left) and incorrect (right) trials across the respiratory cycle. Colored dots indicate circular means, while black arrows represent the direction of the respiratory cycle; 0 rad: exhalation onset; π rad: inhalation onset. The asterisk indicates statistical significance. **C)** ERrS and **D)** rITC assessed for correct and incorrect trials. The gray area represents the window of statistical significance. Red arrows represent the cue onset; the overlaid ECG signal represents an example of an S-trial, included to illustrate the temporal dynamics of the trial with the three target heartbeats highlighted.

### 3.6. Tactile detection task and respiration: accuracy, sensitivity and criterion

TDT accuracy exhibited a trend towards significance, tending to be higher during exhalation than inhalation (t_31_ = -1.95, p = .06, d = -0.35). This trend appears to be driven by a significantly higher false alarm rate during inhalation compared to exhalation (t_31_ = 2.36, p = .025, d = .42, Fig. 4A), while no significant difference was found in the hit rate (t_31_ = -1.69, p = .1, d = -0.3). Task sensitivity (d’) was significantly higher during the exhalation phase compared to the inhalation phase (t_31_ = -6.84, p = 1.17 × 10^−7^, d = -1.21, Fig. 4A), while no significant differences were observed for criterion (c) between inhalation and exhalation (t_31_ = -0.43, p = .67, d = -0.08).

**Figure 4.**
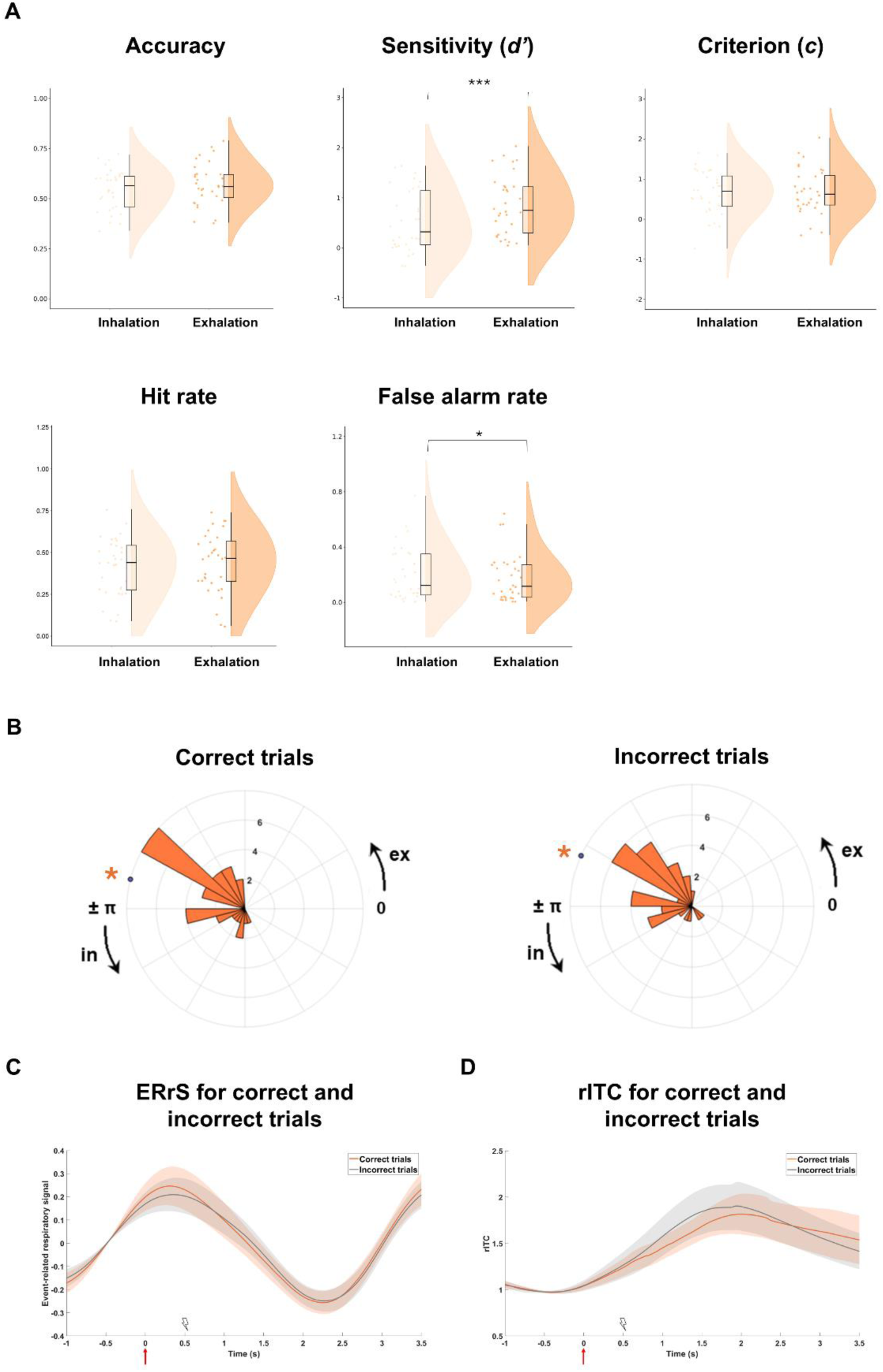
Relationship between respiration and accuracy in the TDT. **A)** Accuracy, sensitivity (d’), criterion (c), hit rate and false alarm rate obtained by dividing the respiratory signal into inhalation and exhalation phases. * p < .05, *** p < .001. **B)** Distribution of correct (left) and incorrect (right) trials across the respiratory cycle. Colored dots indicate circular means, while black arrows represent the direction of the respiratory cycle; 0 rad: exhalation onset; π rad: inhalation onset. The asterisks indicate statistical significance. **C)** ERrS and **D)** rITC assessed for correct and incorrect trials. Gray areas represent the window of statistical significance. Red arrows represent the cue onset; lightning symbol represents the onset of the electrical stimulation.

### 3.7. Generalized linear mixed-effect models

A GLME approach was employed to determine whether the trial-level phase angle of the respiratory signal could predict trial accuracy in both HDT and TDT. The trial-level instantaneous respiratory phase was extracted via Hilbert transform at the time points corresponding to maximum ITC values observed for each task (2.25 seconds after cue onset for HDT and 1.95 seconds for TDT, and modeled using the sine and cosine components. The inclusion of respiratory phase significantly enhanced model fit for both tasks (HDT: LRT = 35.19, p = 2.28 × 10^−8^; TDT: LRT = 15.92, p = .0003). For HDT, respiratory phase had a significant impact on task accuracy, influenced by both the sine (β = 0.009, t = 1.99, p = .046) and cosine components (β = -0.02, t = -5.61, p = 2.16 × 10^−8^). The model also indicated significant main effects of condition (β = -0.19, t = -30.71, p < .0001, Fig. S2A) and confidence (β = 0.13, t = 4.07, p = 4.7 × 10^−5^, Fig. S2B). For the TDT, respiratory phase significantly influenced task accuracy, which was driven by the cosine component (β = -0.02, t = -3.85, p = .0001). The model also showed significant main effects of condition (β = 0.4, t = 68.27, p < .0001, Fig. S2C) and confidence (β = 0.275, t = 8.04, p < .0001; Fig. S2D).

### 3.8. Respiratory phase clustering analyses

To assess the specific direction of the effect found with the GLME, we tested the uniformity of the distribution of phase angles for both correct and incorrect trials across the respiratory cycle with the Rayleigh test. In the HDT, the Rayleigh test indicated a significant clustering of correct trials during exhalation (z = 6.51, p = .001, V_mean_ = 2.60; Fig. 3B-left), whereas the clustering of incorrect trials was not significant (z = 2.50, p = .08, V_mean_ = 2.72; Fig. 3B-right). The Watson-Williams test for equal means between correct and incorrect trials was also not significant (F = .08, p = .78). In the TDT, the Rayleigh test revealed significant clustering of both correct (z = 14.7, p = 6.23 × 10^−8^, V_mean_ = 2.89; Fig. 4B-left) and incorrect trials (z = 14.34, p = 9.98 × 10^−8^, V_mean_ = 2.71; Fig. 4B-right) during exhalation. However, the Watson-Williams test between correct and incorrect trials was not significant (F = .59, p = .45).

### 3.9. Event-related respiratory signal analyses for correct and incorrect trials

To assess whether the effect observed with the GLME approach was driven by respiratory amplitude modulation, we computed the ERrS of correct and incorrect trials in both tasks. In the HDT, ERrS differed significantly between correct and incorrect trials 2.5-3 s after cue onset, with more negative values observed during correct trials (t = -2.8, p = .004, p_FDR_ = .03, d = -0.53; Fig. 3C). In the TDT, no significant differences in breathing patterns between correct and incorrect trials were observed (t = .19, p = .42, d = .001; Fig. 4C).

### 3.10. Inter-trial coherence analyses for correct and incorrect trials

We further investigated whether the observed effect could be associated with respiratory phase consistency by analysing ITC separately for correct and incorrect trials in both tasks. In HDT, ITC values for correct trials significantly increased relative to baseline within a time window of 1 to 3 seconds after cue onset (t = 2.35, p = .013, p_FDR_ = .02, d = .44). For incorrect trials, ITC values increased relative to baseline within a 2 to 3-second time window, though this effect did not survive correction for multiple comparisons (t = 1.79, p = .04, p_FDR_ = .12, d = .34). The comparison between relative ITC values for correct and incorrect trials showed no significant differences (t = .33, p = .39, d = .06, Fig. 3D). In the TDT, for correct trials, ITC values significantly increased relative to baseline within a 1 to 3-second window (t = 4.64, p = 1.99 × 10^−4^, p_FDR_ = 2.99 × 10^−4^, d = .82). A similar effect was observed for incorrect trials, with ITC values increasing within the same time window (t = 3.54, p = 3.99 × 10^−4^, p_FDR_ = 5.98 × 10^−4^, d = .63). The comparison between relative ITC for correct and incorrect trials was not significant (t = .06, p = .47, d = .01, Fig. 4D).

## 4. Discussion

Humans continuously modulate their breathing to align sensory input with specific phases of the respiratory cycle. This mechanism has been associated with faster response times and enhanced perceptual accuracy. In this study, we investigated whether individuals strategically adjust respiration to optimize the processing of anticipated interoceptive and exteroceptive stimuli. We analysed both average respiratory parameters and trial-locked measures, including the event-related respiratory signal (ERrS) and respiratory inter-trial phase coherence (ITC). At the participant level, signal detection theory metrics such as accuracy, sensitivity (d’), and response criterion (c) were computed across tasks and respiratory phases. At the trial level, we employed mixed-effects models to examine whether the respiratory phase predicted perceptual accuracy. Additionally, by categorizing trials as correct or incorrect, we tested whether these outcomes were preferentially distributed across specific phases of the respiratory cycle. Finally, we assessed whether trial-locked respiratory measures differed between correct and incorrect responses in both tasks. Based on previous literature (Grund et al., 2022; Zaccaro et al., 2022, 2024), we hypothesized that participants would exhibit respiratory modulation during both tasks, aligning exhalation with trial onset to enhance perceptual accuracy.

Results indicated that participants modulated their average respiratory parameters across tasks, exhibiting a significantly lower breathing frequency during the HDT compared to the TDT. Analyses of the ERrS further revealed task-specific respiratory patterns time-locked to trial onset, with lower-amplitude respiration observed during the HDT relative to the TDT. Participants exhibited respiratory phase synchronization to trial onset in both tasks. This was reflected in a significant increase in respiratory ITC, emerging approximately 1.5 seconds after cue onset in the HDT and around 1 second in the TDT, relative to baseline.

Signal detection theory analyses showed that, during the HDT, both accuracy and hit rate were higher during exhalation compared to inhalation. In the TDT, sensitivity was higher during exhalation, accompanied by a lower false alarm rate in the same respiratory phase. These findings are consistent with previous studies that have demonstrated enhanced neural representation of cardiac signals, as indicated by increased HEP amplitude and improved interoceptive accuracy, during exhalation (Zaccaro et al., 2022, 2024). In the exteroceptive domain, our results replicate those reported by Grund et al. (2022) using a similar tactile detection paradigm and are in line with a broader body of evidence showing increased sensory responsiveness during exhalation. This includes increased startle responses to auditory stimuli (Münch et al., 2019), stronger conditioned learning (Waselius et al., 2019, 2022), better audio-tactile and audio-visual multisensory integration (Saltafossi et al., 2025), and greater auditory-evoked potentials (Mizuhara et al., 2025) and mismatch negativity amplitudes (Mizuhara et al., 2024).

At the trial level, separate mixed-effects models for the HDT and TDT showed that including the respiratory phase angle as a predictor for accuracy significantly improved model fit. This suggests that the ongoing respiratory phase during task performance plays an important role in modulating perceptual accuracy in both tasks. We also tested the distribution of the trial-level respiratory phase angles across the respiratory cycle separately for correct and incorrect trials. During the HDT, correct trials were significantly clustered around the end of the exhalation phase, whereas incorrect trials showed no significant phase clustering. In contrast, during the TDT, phase angles were significantly clustered around the end of exhalation for both correct and incorrect trials. We then compared trial-locked respiratory activity between correct and incorrect trials in both tasks. In the HDT, ERrS amplitude was significantly more negative between 2.5 and 3 seconds after cue onset for correct compared to incorrect responses. Since the ERrS had a negative value, we suggest that, on average, participants were exhaling during correct trials, whereas incorrect trials were more likely to occur during inhalation. In the TDT, no significant ERrS differences between correct and incorrect trials emerged. Finally, during the HDT, we found increased respiratory ITC relative to baseline specifically for correct trials, but no significant difference emerged when comparing correct and incorrect trials. In contrast, during the TDT, ITC was higher relative to baseline for both correct and incorrect trials, with no significant difference between conditions. These findings underline a link between respiratory phase alignment and task performance in the interoceptive domain while suggesting that phase-locking in the exteroceptive domain is less strongly associated with perceptual accuracy.

Taken together, these results confirm that specific phases of the respiratory cycle facilitate the perception of near-threshold stimuli in both the interoceptive and exteroceptive domains. Moreover, they support the view that respiration is not merely a passive modulator but is actively regulated by individuals to optimize perceptual outcomes, consistent with the interoceptive predictive coding framework (Barrett & Simmons, 2015; Seth, 2013; Seth & Friston, 2016). Specifically, participants appeared to align expected sensory events with preferred phases of the respiratory cycle. Although this behaviour has been previously demonstrated in the exteroceptive domain (Grund et al., 2022; Harting et al., 2025; Huijbers et al., 2014; Johannknecht & Kayser, 2022; Kluger et al., 2021; Perl et al., 2019), our findings extend this mechanism to the cardiac interoceptive domain, providing support for the notion of a respiratory active inference strategy (Boyadzhieva & Kayhan, 2021; Corcoran et al., 2018): individuals appear to modulate their respiratorion (both in phase and amplitude) to enhance the perception of internal bodily signals. We refer to this process as interoceptive respiratory active sensing.

We propose that, although the phase-locking of exhalation to sensory inputs appears behaviourally similar across tasks, it is driven by distinct underlying mechanisms shaped by specific task demands. In the HDT, respiratory phase modulates cardiac interoception through a set of hierarchically organized processes (Smith et al., 2017). At the physiological level, exhalation enhances baroreceptor signalling due to respiratory sinus arrhythmia (Brecher & Hubay, 1955; Larsson et al., 2021). At the subcortical level (mostly involving vagal pathways and the nucleus of the solitary tract) respiratory-related afferents may induce interoceptive noise that masks cardiac signals when systoles occur during inhalation (Noble & Hochman, 2019; Streeter et al., 2012). This is consistent with findings showing improved interoceptive accuracy during breath-holding conditions, where respiratory interference is minimized (Lavalley et al., 2024; Smith et al., 2020, 2021). At higher cortical levels, exhalation is associated with reduced cortical excitability, which may facilitate interoceptive attention due to the suppression of task-irrelevant distractors. This is reflected in increased heartbeat-evoked potentials, as well as heartbeat-related alpha power and ITC (Zaccaro et al., 2022, 2024, 2025). Consequently, individuals may initiate exhalation to actively increase the perceptual salience of heartbeat sensations (Ainley et al., 2016; Larsson et al., 2021; Leganes-Fonteneau et al., 2021; Molle & Coste, 2022).

In contrast, during the TDT, the role of exhalation appears less straightforward. In the auditory domain, enhanced sensory responses during exhalation have been attributed to a release of cognitive resources, since exhalation inherently demands less neural engagement than inhalation (Feldman & Del Negro, 2006; Muñoz-Ortiz et al., 2019). These available resources may then be reallocated to facilitate the processing of weak or ambiguous sensory inputs (Mizuhara et al., 2024, 2025). An alternative explanation, grounded in the predictive coding framework, suggests that in tasks requiring precise somatosensory perception, such as the TDT, exhalation may be actively initiated to induce a parasympathetically mediated reduction in heart rate that, rather than enhancing the salience of cardiac signals. This may in turn minimize baroreceptor-driven interference, which could disrupt the temporal and spatial processing of tactile stimuli (Al et al., 2020, 2021; Grund et al., 2022; Saltafossi et al., 2023). This account is supported by evidence showing that cardiac deceleration often precedes the presentation of salient exteroceptive stimuli, possibly serving as a preparatory mechanism to enhance sensory processing (Skora et al., 2022).

While this study offers important contributions, it is important to note some limitations. First, HDT is not a purely interoceptive measure, as it requires integration of both interoceptive and exteroceptive signals. Nevertheless, HDT performance is less susceptible to confounding factors such as prior knowledge of one’s resting heart rate or time estimation abilities (Phillips et al., 1999; Tanaka et al., 2021). Second, the use of fixed time windows for S+ and S– trials may have reduced task sensitivity due to interindividual variability in cardiac signal perception (Lyyra & Parviainen, 2018; Schulz et al., 2013, 2021; Sokol-Hessner et al., 2015; Wiens et al., 2000). Adopting a modified version of the paradigm, such as the method of constant stimuli, in which auditory tones are presented at multiple latencies following the R-peak, could improve sensitivity and generalizability of results (Brener et al., 1994; Clemens, 1984; Mailloux & Brener, 2002; Wiens & Palmer, 2001; Yates et al., 1985). Finally, as this study did not employ EEG, any interpretation of respiration in modulating the higher-order neural basis of cardiac and somatosensory perception remains speculative. Future EEG research could help clarify the neural mechanisms underlying respiratory-phase modulation of both interoceptive and exteroceptive accuracy. A more comprehensive understanding of how breathing actively contributes to shaping internal and external perception may support the development of novel therapeutic strategies that target respiratory control to enhance interoceptive and exteroceptive functioning.

## Acknowledgments

This research paper was supported by Boosting Ingenium for Excellence (BI4E) project, funded by the European Union’s HORIZON-WIDERA-2021-ACCESS-05-01-European Excellence Initiative under the Grant Agreement No. 101071321; the National Recovery and Resilience Plan (NRRP), Mission 4, Component 2, Investment 1.1, Call for tender No. 1409 published on 14.9.2022 by the Italian Ministry of University and Research (MUR), funded by the European Union - NextGenerationEU - Project Title “Metaphor end epistemic injustice in mental illness: the case of schizophrenia” - CUP D53D23020890001- Grant Assignment Decree No. 1409/2022 adopted on October 31 2023, by the Italian Ministry of University and Research (MUR); the Departments of Excellence 2023–2027” initiative of the Italian Ministry of University and Research for the Department of Neuroscience, Imaging and Clinical Sciences (DNISC) of the University of Chieti - Pescara; the National Recovery and Resilience Plan (NRRP), Mission 4, Component 2, Investment 1.1, Call for tender No. 104 published on 2.2.2022 by the Italian Ministry of University and Research (MUR), funded by the European Union - NextGenerationEU - Project Title “Interception and Active Aging (InterActing)” - CUP D53D23009700006 - Grant Assignment Decree No. 1016 adopted on July 7 2023 by the Italian Ministry of University and Research (MUR).

## Declaration of interests

The authors declare no competing interests.

## Author contributions

**Francesca della Penna:** conceptualization, methodology, software, validation, formal analyses, investigation, data curation, visualization, writing-original draft, writing-review & editing. **Andrea Zaccaro**: conceptualization, methodology, software, formal analyses, data curation, writing-original draft, writing-review & editing. M**auro Gianni Perrucci**: methodology, software, data curation, resources, writing-review & editing. **Marcello Costantini**: visualization, supervision, funding acquisition, writing-review & editing. **Francesca Ferri**: conceptualization, methodology, resources, supervision, project administration, funding acquisition, visualization, writing-review & editing.

## Notes

### Competing Interest Statement

The authors have declared no competing interest.

