## Supplemental material for "Breathing strategies to influence perception: Evidence for interoceptive and exteroceptive active sensing"

**Pilot study**

*Methods.* Thirty healthy volunteers (20 females; 1 left-handed; age: 26.2 ± 5.1 years [mean ± SD]) were recruited to complete three blocks of a modified heartbeat discrimination task (HDT), with each block corresponding to a different condition: 3 tones, 2 tones, and 1 tone. Each block contained 50 trials, for a total of 150 trials, with an equal distribution of simultaneous (S+) and non-simultaneous (S-) trials. In the S+ condition, the tone onset was delayed by 250 ms after the participant’s R-peak onset, while in the S- condition, the tone onset was delayed by 550 ms after the R-peak onset. After each trial, participants were asked to decide whether they perceived the heartbeat and the tones as simultaneous by pressing a specific key on the computer keyboard, and to rate their confidence on a 4-points Likert scale, ranging from 1 (very unconfident) to 4 (very confident).

The HDT was preceded by an exteroceptive audio-visual familiarization task (AV Familiarization task). In the AV Familiarization task (Betka et al., 2021), participants were required to judge the simultaneity between a flashing dot and a sequence of acoustic signals played through computer speakers. The goal of this task was to familiarize participants with the judgement of simultaneity. There were two conditions: simultaneous (S+) and non-simultaneous (S-). In the S+ condition, the flashing dot and tone were presented without delay, while in the S- condition, the tone delivery onset was delayed by 550 ms after the flashing dot onset. The task consisted of 3 blocks corresponding to three different conditions: 3 tones, 2 tones, and 1 tone. Each block contained 10 trials with an equal distribution of S+ and S- trials, presented in random order. After each trial, participants had to decide whether they perceived the flashing dot and tones as simultaneous or not by pressing a specific key on the computer keyboard. They also rated their confidence on a 4-point Likert scale, ranging from 1 (very unconfident) to 4 (very confident).

Cardiac and respiratory activity were simultaneously recorded using three Ag/AgCl pre-gelled electrodes placed over the left and right clavicles and the right costal margin (ground electrode), along with a respiratory belt placed over the chest (respiratory transducer TSD201, BIOPAC Systems, Inc).

*Results.* The results of the Pilot study showed that, in the 1-tone block, only 23% of participants achieved a performance with accuracy greater than 55% (indicating significantly above chance levels), compared to 47% in the 3-tone block, while in the 2-tone condition 43% of participants achieved an acceptable performance. These results are consistent with those reported by Brener in 1994, in which he compared the use of different numbers of consecutive tones (1, 5, and 10, Brener et al., 1994). According to Brener, the presentation of a single tone is insufficient to obtain a reliable measure of cardiac interoceptive accuracy, as only 13% of participants were able to perceive the heartbeat with 1 tone, compared to 47% with 10 tones. Brener suggested that the higher proportion of participants able to perceive the heartbeat with multiple tones is due to the processes of orientation and attention facilitated by the sequence of stimuli. When participants have difficulty perceiving their heartbeat, the sequence of tones provides a temporal window in which they can locate the heartbeat, aiding its perception. The interval between tones likely provides a period that allows participants to anticipate the next heartbeat (Brener et al., 1994). Based on our results and those from previous studies, we decided to proceed with the 3-tones condition. Although there remains a risk that consecutive heartbeats may span multiple respiratory phases, this approach ensures that more than 50% of the heartbeats in each trial (i.e., two out of three) fall within the same respiratory phase, allowing us to categorize the entire trial as occurring either during inhalation or exhalation.

To make sure that the observed effects on Event-related respiratory Signal in the 3-tone condition were not caused by the presentation of three consecutive stimuli instead of one, as in the exteroceptive condition, we evaluated the Event-related respiratory Signal in the 1-tone condition of the Pilot study and compared it to the Event-related respiratory Signal assessed for the TDT (Fig. S1); since only 16 (12 females; age: 27 ± 6.39 years [mean ± SD]) out of 30 participants in the Pilot study met the inclusion criteria (accuracy above 55% and d' > 0) in the 1-tone condition, we reduced the exteroceptive sample to 16 participants (7 females; 1 left-handed; age: 25.25 ± 5.2 years [mean ± SD]) to ensure a more accurate comparison. However, it is important to note that the number of trials differed consistently between the two conditions (50 trials in the 1-tone condition of the Pilot study compared to 300 trials in the TDT). The comparison revealed a significant difference between 0 and 1 second after cue onset (t = 2.21, p = .01, p_FDR_ = .04, d = .14), indicating that the respiratory modulation is not driven by the number of stimuli, but rather reflects a consistent strategy adopted across conditions.

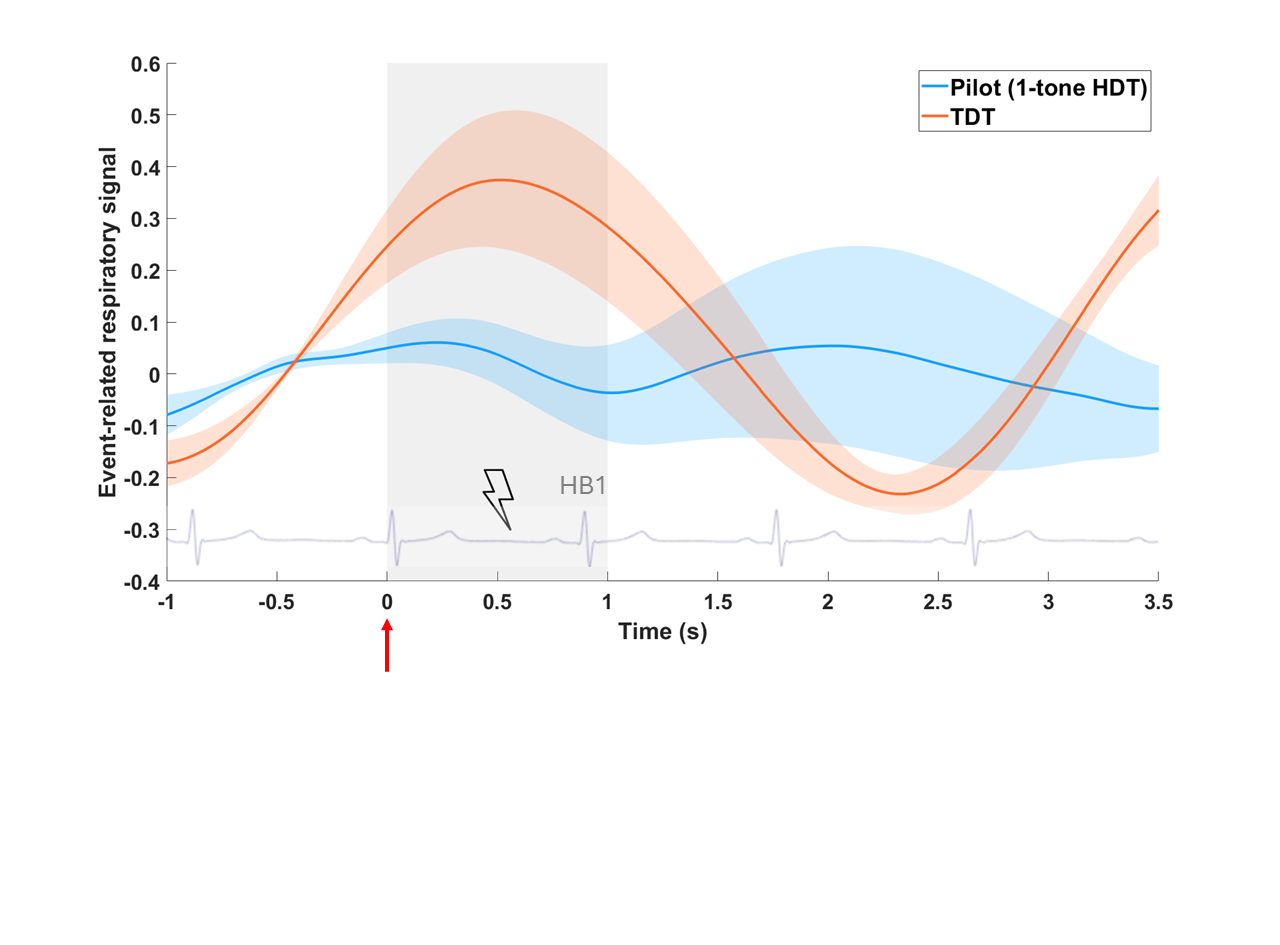

**Figure S1**. Event-related respiratory signals assessed for the 1-tone condition of the Pilot study and the exteroceptive task (TDT). The gray area represents the window of statistical significance. The red arrow represents the cue onset in both conditions; lightning symbol represents the onset of the electrical stimulation in the TDT; the overlaid ECG signal represents an example of an S- trial, included to illustrate the temporal dynamics of the trial with the target heartbeat highlighted.

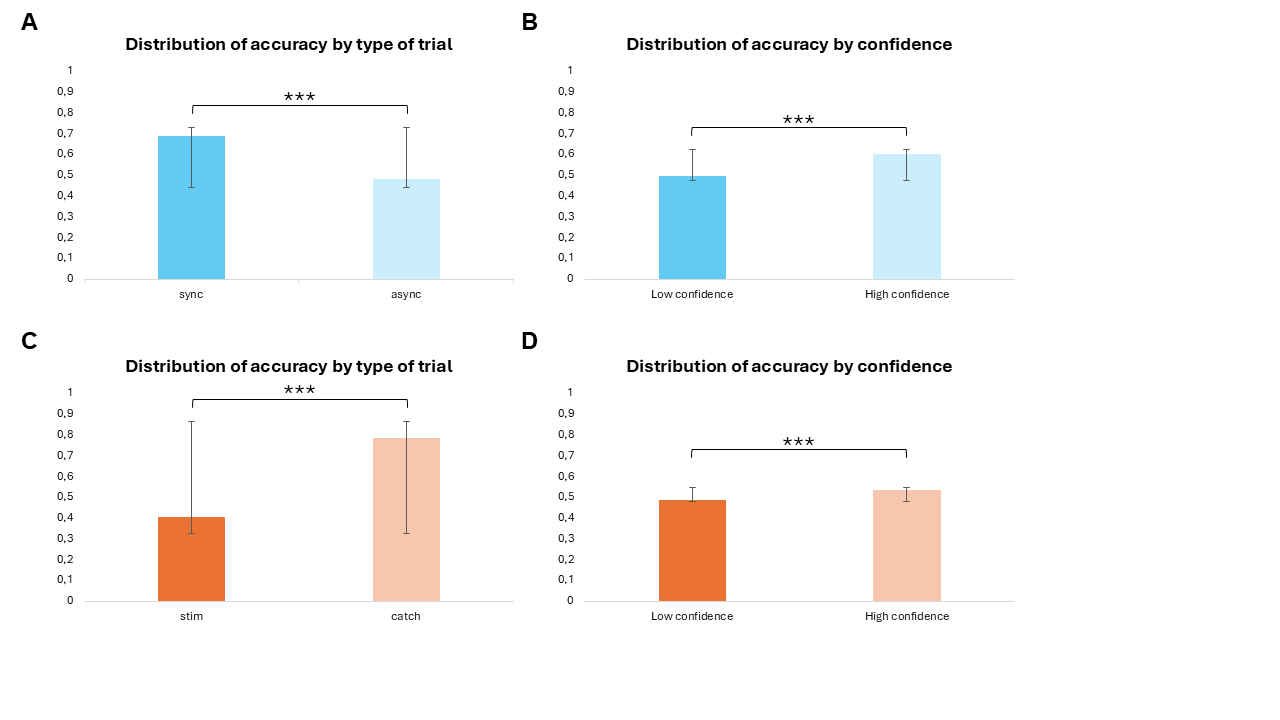

**Figure S2.** Distribution of accuracy by type of trial (A, C) and confidence (B, D) in the HDT (top) and TDT (bottom). *** p < .001.
